# Differential Neuroanatomical, Neurochemical, and Behavioral Impacts of Early-Age Isolation in a Eusocial Insect

**DOI:** 10.1101/2023.06.29.546928

**Authors:** Billie C. Goolsby, E. Jordan Smith, Isabella B. Muratore, Zach N. Coto, Mario L. Muscedere, James F. A. Traniello

## Abstract

Social experience early in life appears to be necessary for the development of species-typical behavior. Although isolation during critical periods of maturation has been shown to impact behavior by altering gene expression and brain development in invertebrates and vertebrates, workers of some ant species appear resilient to social deprivation and other neurobiological challenges that occur during senescence or due to loss of sensory input. It is unclear if and to what degree neuroanatomy, neurochemistry, and behavior will show deficiencies if social experience in the early adult life of worker ants is compromised. We reared newly-eclosed adult workers of *Camponotus floridanus* under conditions of social isolation for 2 to 53 days, quantified brain compartment volumes, recorded biogenic amine levels in individual brains, and evaluated movement and behavioral performance to compare the neuroanatomy, neurochemistry, brood-care behavior, and foraging (predatory behavior) of isolated workers with that of workers experiencing natural social contact after adult eclosion. We found that the volume of the antennal lobe, which processes olfactory inputs, was significantly reduced in workers isolated for an average of 40 days, whereas the size of the mushroom bodies, centers of higher-order sensory processing, increased after eclosion and was not significantly different from controls. Titers of the neuromodulators serotonin, dopamine, and octopamine remained stable and were not significantly different in isolation treatments and controls. Brood care, predation, and overall movement were reduced in workers lacking social contact early in life. These results suggest that the behavioral development of isolated workers of *C. floridanus* is specifically impacted by a reduction in the size of the antennal lobe. Task performance and locomotor ability therefore appear to be sensitive to a loss of social contact through a reduction of olfactory processing ability rather than change in the size of the mushroom bodies, which serve important functions in learning and memory, or the central complex, which controls movement.

Visual Abstract.
Newly eclosed (callow) *Camponotus floridanus* minor workers were subjected to variable periods of isolation to determine whether and how diminished social experience and isolation influence the development of the brain, including compartment volumes, biogenic amine levels, and behavioral performance.

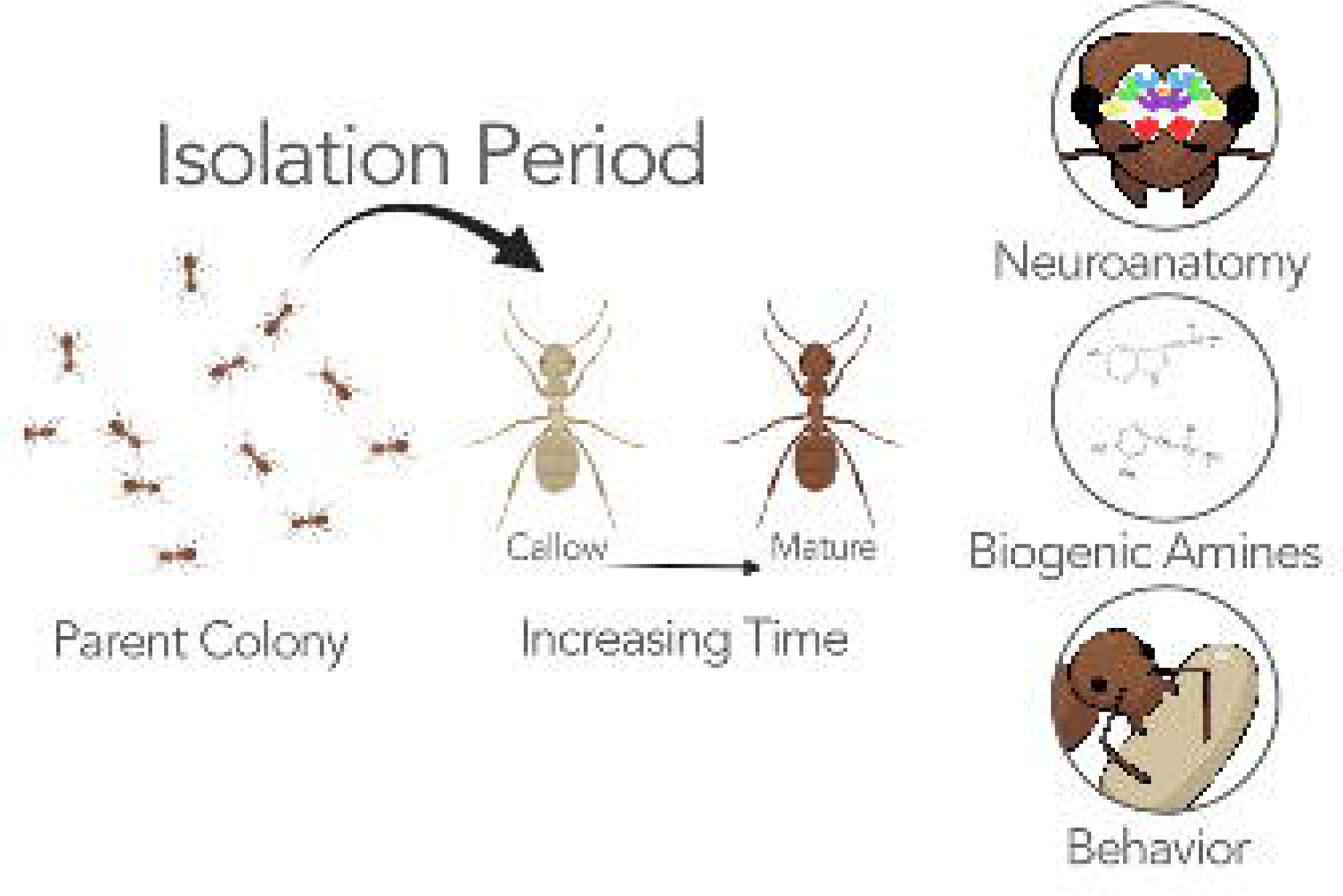

## Introduction

Social experience shapes the nervous system and the behavior it generates. Exposure to social stimuli and cues is critically important to neural development, behavioral performance and cognition [1–5], neuroimmune function [6–10], and most significantly, survival [11, 12]. Across diverse clades, behavioral capabilities can be negatively impacted if individuals are deprived of social contact when ontogenic trajectories are most sensitive to receiving sensory inputs [13, 14]. A lack of social contact early in life can disrupt maturational processes required for feeding, aggression, courtship, and interaction networks [12, 15].

Although research on the effects of social isolation has historically focused on humans, non-human primates, and other mammals, eusocial insects have emerged as important systems to examine the influences of social deprivation in early adult life on brain development, brain gene expression, neuromodulators, and behavior [16–23]. Workers and immatures are developmentally dependent [24]. Social interactions such as trophallaxis (oral food exchange), allogrooming, and nursing influence social network structure and behavioral and neuroanatomical development trajectories [23, 25–27]. Age-related changes in functionally specialized brain compartments can be experience-expectant (growth programmed to occur despite experiential input) or experience-dependent growth resulting from the performance of a particular activity, such as foraging [28, 29]. Social isolation in ants appears to significantly alter the growth of the mushroom body, a brain compartment involved in higher-order sensory processing, learning, and memory, relative to that of nestmates developing in a natural social environment [17]. If workers are deprived of social contact, neuromodulators and receptor profiles that regulate behavior may be altered or become dysregulated [16, 30], potentially affecting age-related responsiveness to signals and cues that guide neural and behavioral development, and task performance inside and outside of the nest [20, 31–37]. Nevertheless, worker brains and behavior may be resilient to senescence [38] and injury to sensory appendages [39]. This robustness to aging and loss of olfactory input from the antennae raises questions concerning the ability to compensate for stressors resulting from social isolation during early adulthood and whether worker neurobiology and behavior are equally sensitive to such disruption.

In ants, newly eclosed adult workers (callows) typically develop in a socially rich colony environment of demographically variable nestmates and immatures. Callows are distinguished from their mature siblings by their unsclerotized lightly pigmented cuticles and relatively low activity [40, 41]. Behavioral repertoire formation is usually incomplete immediately following adult eclosion, with apparently few exceptions [42]. Manipulations of the social experience of callows therefore provides the ability to experimentally examine the ontogeny of behavior from the earliest stage of adult life. Social interaction between callows and their siblings influences pheromone responsiveness and temporal behavioral development [40, 43–46]. The effects of being deprived of these and other inputs are not consistently negative: 10-day old *Pheidole dentata* minor workers that lacked contact with brood nursed as efficiently as same-aged control workers and did not differ in brain anatomy or neurochemistry [20]. The question remains as to whether neuroanatomy, neuromodulators, behavioral responsiveness to social signals (such as those regulating brood care), and stimuli involved in sensing the extranidal environment (such as those involved in foraging) develop independently of worker interactions with colony members early in life and are differentially influenced by social deprivation.

The ant genus *Camponotus* has served as a model to examine neuroanatomical plasticity during development [28] and neuroethological and neurochemical impacts of social deprivation. In *C. floridanus* and *C. fellah*, decreases in worker mushroom body size scaling, changes in levels of biogenic amines, and deficits in social behavior have been documented in isolated individuals [16, 17, 30, 47]. Here, we integrate analyses of brain size and the growth of neuropils specialized for sensorimotor functions, titers of brain biogenic amine neuromodulators, and task performance competence to examine how a lack of social contact during early life affects worker neurobiology and behavior.

## Methods

### Colony collection and culture

Three queenright colonies of *C. floridanus* were collected in or near Gainesville, Florida in 2012 and reared in Harris environmental chambers at 25°C and 40-55% relative humidity on a 12:12 light cycle. Parent colonies from which workers were sampled had a minimum size of 100, equally distributed between minors and majors. Artificial nests were made from test tubes 75% filled with water and fitted with a tight cotton plug and placed in Fluon®-lined (Bioquip, Rancho Dominguez, CA) plastic nest boxes 16×11×6 cm to 32×22×6 cm depending on colony size. Colonies were fed 1M sucrose on saturated cotton balls and varied protein sources (mealworms, scrambled eggs, and fruit flies) every other day in rotation.

### Worker social isolation

*C. floridanus* has a dimorphic worker caste. We used minor workers because of their diverse behavioral repertoire, which includes brood care and foraging, rather than majors, which are limited to defense. Four groups of callow minors were collected from each of three parent colonies. Randomly selected callows were isolated from nestmates for 2-3 days (n=29), 5-6 days (n=36), 20-22 days (n=44), and 40 days (n=29, mean = 40.08 days, range = 30 – 53 days; herein 40 days) in separate Fluon®-lined boxes (16 × 11 × 6 cm). Socially isolated workers (“isolates”) were provided similar conditions in smaller nest boxes and the same diet as parent colonies. After the prescribed isolation period, isolates were assayed for behavioral performance and subsequently sacrificed for immunohistochemical analysis of brain structure, or for quantification of brain biogenic amine titers. Callow (n = 29) and mature minor workers (“matures”, n = 46) were collected and assayed using the same methods. Callow and mature controls served as comparative groups for 2-day isolates and 40-day isolates, respectively, and we report primarily on differences in behavior, neuroanatomy, and neurochemistry between these groups. We compared brain volumes of callow controls estimated to be 2 days old and 2--3-day isolates to ensure there were no intrinsic differences in brain size post eclosion. We used 5- and 20-day isolates lacking age-matched controls to estimate the trajectory of behavioral and neural development from 2 days to 53 days of isolation (**Supp Fig 1-4**). Isolated workers had a survival rate of 63.7%. Because of the relatively high mortality of workers experiencing longer periods of isolation, sample sizes for 40-day isolates are generally lower. Head width, measured as the widest distance across the eyes, served as a proxy for body size. To control for the effect of body size on brain size, we compared the head widths of workers across isolation periods and control groups and found no significant differences (p=0.40, R^2^=0.098, F=1.18, Z=0.25, SS type 3, **Supp Fig 1a)**. Our sampling of workers for assignment to isolation or control treatments was thus unlikely to influence brain volume.

### Behavioral performance assays

Isolated and control workers used for behavior (n = 16 [2 day], 19 [5 day], 20 [20 day], 11 [40 day], 8 [callow] and 13 [mature]) were exposed to a randomized series of assays to determine foraging ability and ability to recognize and care for brood, tasks that affect colony fitness. Simplified illustrations of the assays can be found in **Fig 1a**.

**Figure 1.**
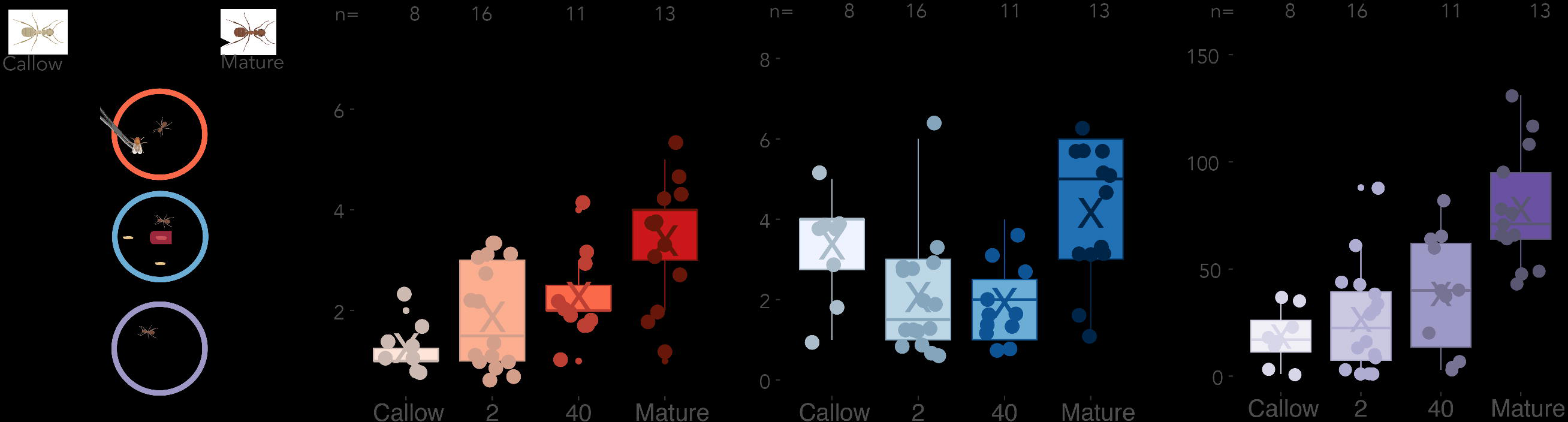
Isolation effects on task performance. **(a)** Illustration of brood-care and predatory behavior assays. **(b)** Predation scores of workers isolated for varying time periods compared to control callow and mature workers. **(c)** Brood-care scores of isolates compared to callows and mature worker controls. **(d)** Locomotion scores of isolates, callows and matures. X in box plot = average; horizontal line = median. Sample sizes are above significance indicators. <0.001 = ***; 0.001-0.01 = **; 0.01-0.05 = *; >0.05 = ns.

#### Predatory Behavior

A single worker was carefully placed in a Fluon®-lined Petri dish (9cm) and allowed to acclimate for 2 minutes prior to assay initiation. Using Dumont No. 5 fine forceps sanitized with ethanol and dried, a live fruit fly was offered to the worker and the response was observed for 2 minutes. The degree of predatory behavior was scored on a 5-point scale; higher scores indicate greater predatory behavior. 1. no response or avoidance; 2. olfactory (antennal) investigation; 3. mandibular flaring; 4. latent attack (delayed and intermittent); and 5. immediate attack (persistent predatory attack from first encounter with prey). The assay arena was also sanitized with ethanol between trials to minimize influence from social signals or prey odors.

#### Brood recognition and care

To determine if isolates differ in the ability to recognize and care for immature siblings with similar efficacy as control matures and callows, we evaluated their social interactions with pupae. Each focal worker was carefully placed in a 14 cm diameter Petri dish test arena where a 30 × 10 mm red plastic hollow tube provided a darkened area into which workers could move pupae. Three pupae were added at three different locations equidistant from the darkened chamber before a worker was introduced. Workers could relocate pupae either inside or outside of the dark chamber or have no interaction. Pupae were recorded as aggregated (“piled”) if workers moved two or more together at any location. Workers showing this level of response began pupal care immediately. We did not assign lower scores to workers that clustered pupae without moving them into the darkened area. The darkened tube provided the opportunity for a level of pupal care that would emulate natural placement of immatures within a colony.

However, no worker moved pupae into the tube and we therefore did not include this behavior in the calculation of brood-care scores. Some workers did not move any pupae, but they themselves moved into the darkened tube. This was scored as avoidance.

Worker interactions with pupae were observed directly for 5 minutes; after an additional 15 minutes, the final placement of pupae was recorded (total trial length = 20 minutes). Brood recognition and care was measured on a 7-point scale: 1. no interaction or avoidance throughout the duration of the trial; 2. olfactory (antennal) investigation; 3. mouthpart contact; 4. one pupa moved during the last 15-minute period; 5. one pupa moved in the first five-minute period; 6. two pupae moved beginning in the first five-minute period; 7. three pupae moved beginning in the first five-minute period. Levels 4 and 5 distinguish the rate of response to pupae, which we interpret as sensory processing ability that could be sensitive to social isolation. Higher scores indicate greater brood-recognition ability (likely olfactory), restoration of brood clusters through pupal transport, and sustained engagement with brood. Workers received scores of 5-7 independent of where pupae were transported.

#### Locomotion

To examine if isolation affected physiological and/or biomechanical capability to move pupae or attack prey, we used a simple and robust proxy to broadly assess neuromuscular impairment [38]. Focal workers were placed in the center of a 10 cm diameter Fluon®-lined Petri dish divided into four equal quadrants by a crosshair pattern drawn beneath. After a two-minute acclimation period, the number of times a worker moved among the four quadrants was recorded during a five-minute period using a digital hand counter. Movement between quadrants was only scored when the worker passed a division line by at least one body length.

### Neuroanatomical and neurochemical measurements

#### Immunohistochemistry and confocal microscopy

Intact brains (n = 8 [2 day], n = 11 [5 day], n = 15 [20 day], n = 14 [40 day], n = 5 [matures], and n = 7 [callows]) were dissected in ice-cold Ringer’s solution, and then immediately placed in ice-cold zinc-formaldehyde fixative with shaking overnight. Fixed brains were then washed in HEPES buffered saline 6 times (10 minutes/wash) and fixed in Dent’s Fixative (80% MeOH, 20% DMSO) for 1 hour. Brains were placed in 100% methanol and stored at −17 C° for 1-3 weeks until processed. Brains were next rinsed 6 times (10 minutes/rinse) with 3% Triton X-100 in 1X PBS (3% PBST). Brains were stored overnight in 5% normal goat serum/0.2% PBST/0.005% sodium azide for blocking. After blocking, brains were incubated for 4 days RT with an anti-synapsin antibody to stain neuropils (1:30, Development Studies Hybridoma Bank, Iowa City, Iowa) in constant nutation. Brains were washed with 0.2% PBST (6×10 minutes), then incubated in secondary antibody (1:100 AlexaFluor goat L-mouse) in 5% normal goat serum/0.2%PBST/0.005% sodium azide for 4 days at RT with constant nutation. Brains were washed in 0.2% PBST (6×10 minutes), then dehydrated through an ethanol and PBS series (30, 50, 70, 95, 100% ethanol in 1XPBS). Brains were cleared and immersed in methyl salicylate and mounted on stainless steel glass-windowed slides for confocal imaging.

Brains were imaged with a Nikon C2 + Si spectral laser scanning confocal microscope (Nikon, Melville, NY, USA) with sections at approximately 1.5 um throughout the entire brain. Images were manually annotated using Amira 6.0 software to quantify neuropil volumes. The individual who annotated brains was blind to brain sample category, thus minimizing human bias. The margins of neuropils were identified visually or with magic wand tools in Amira. Every eighth frame was annotated manually. Biomedisa interpolation [48] was used on frames without labels, then evaluated and edited by the annotator for accuracy. We labeled volumes of the optic lobe (OL; vision), antennal lobe (AL; olfaction), mushroom body (MB; higher-order processing, learning, and memory), lateral calyx (MB-LC), and medial calyx (MB-MC), peduncles (MB-P), central complex (CX; movement and navigation), suboesophageal zone (SEZ; mouthpart control), and rest of the central brain (ROCB). ROCB describes tissue composed of the superior neuropils, ventromedial neuropils, ventrolateral neuropils, and the lateral horn [49]. For each brain compartment, volumes of only one intact hemisphere were recorded. If both hemispheres were intact, the labeler chose one randomly.

A detailed description of brain anatomy and development in relation to age and experience in *C. floridanus* is presented in Gronenberg et al. [28]. We compared the relative volumes of brains across all groups and scaling relationships among brain compartments. Relative volumes were calculated by dividing the volume of each focal compartment by the ROCB volume. Damaged compartments were excluded from analysis. If one or more compartments or the ROCB was damaged, the sample was excluded. The MB-LC and MB-MC exhibited similar volumetric trends and thus were combined to one metric (MB-C).

#### Quantification of brain biogenic amines

Isolated workers were quickly dissected (n = 11 [2 days], n = 10 [5 days], n = 12 [20 days], n = 3 [40 days], along with control mature workers (n = 11) to quantify levels of serotonin (5HT), dopamine (DA), and octopamine (OA) using isocratic, reversed-phase high-performance liquid chromatography with electrochemical detection (HPLC-ED). Monoamine quantification was performed blind to treatment. The HPLC-ED system (formerly ESA, Inc., Chelmsford, MA, USA) included a model 584 pump, a model MD-150 (3 × 150 mm) reversed-phase analytical column, a model 5011A dual-channel coulometric analytical cell, and a Coulochem III electrochemical detector. Brains were dissected in less than 5 minutes in ice-cold Ringer’s solution, homogenized in mobile phase manually with a plastic polypropylene pestle in a centrifuge tube, and centrifuged. The supernatant was then injected into the HPLC column, and monoamine levels were quantified with reference to external standards. Mobile phase was formulated as 50 mM citrate/acetate buffer, 1.5 mM sodium dodecyl sulfate, 0.01% triethylamine, and 25% acetonitrile in MilliQ water [20]. Multiple daily runs of the external standards accounted for any changes in ambient conditions.

### Statistical analysis and data visualization

All statistics analyses were performed in RStudio version 4.2.3 [50]. For generalized linear mixed models, we used the package ‘*RRPP’* [51]. We tested for statistical significance using ANOVA with residual randomization in a permutation procedure of 1000 iterations and the estimation method of ordinary least squares, given combined sample size of 25 or more. We included colony and head width as random effects in our behavior, brain volume, and biogenic amine modeling; neither had a significant effect. Isolation significantly correlated with head width only when head width was nested with the microscope and experimenter, rather than head width alone, meaning that slight variation between ocular micrometers of the microscopes, rather than treatment, influenced the effect of head width. ANOVA and modeling details can be found in **Supp Tables Set 1 and 2.** Following ANOVA, *post-hoc* tests were done on pairwise comparisons using Wilcoxon rank sum exact tests with a Benjamini-Hochberg (BH) procedure to correct for multiple tests. Depending on the sample size of 40-day isolates, we included type III error to account for sample size bias. Additional nonparametric statistical analyses including Kruskal-Wallis H tests and *post-hoc* pairwise comparisons using t tests with a Bonferroni correction, which agreed with our modeling, can be found in **Supp Tables Set 1 and 2.** We then ran studentized Breusch-Pagan (BP) tests from the ‘*lmtest’* package [52] to confirm homoscedasticity in our model. If heteroscedasticity was found in raw values, we log transformed our data and reran the model and confirmed homoscedasticity. Heteroscedasticity could not be removed from only our mixed model analysis of MB-P scaled volumes in **Supplementary Tables, Set 1**. However, the model nonetheless agrees with our additional nonparametric statistical tests. We confirmed that log-transformed data resulted in the same pairwise comparison results as raw data, confirming that log transformation for model fitness did not create false pairwise findings. Because 5-day isolates and 20-day isolates did not have socially typical, age-matched controls and were instead meant to provide insight into the trajectory of neurodevelopment and behavior, we excluded these groups from the analysis reported in the manuscript. Our analyses including these excluded groups can be found in **Supp Figs 1-4** and in **Supp Tables, Set 2.** Figures were created using the package ‘*ggplot2*’ [53].

Figure assembly was done in Adobe Illustrator (Adobe Illustrator 2023). Illustrations of ants in **Visual Abstract, Fig 1a**, and **Supp Fig 1b** are courtesy of BioRender.com.

## Results

### Effect of Isolation on Predatory Behavior

Mature control workers preyed on fruit flies significantly more (x□ = 3.39) than workers in all other experimental groups and callows (vs. callows, 2-day isolates, and 40-day isolates: p < 0.029; BH correction). Forty-day isolates exhibited significantly higher predatory behavior (x□ = 2.27) than callows (x□ =1.25; p=0.0093, BH correction). All other pairwise comparisons were not significantly different (p=0.16-0.29). **(Fig 1a, b)**. Therefore, control callows and 2-day isolates similarly avoided prey or showed olfactory investigation whereas older isolates (40-day, e.g.) had higher predatory scores that resembled those of mature worker controls but nevertheless were significantly lower.

### Effect of Isolation on Brood Recognition and Care

Brood-care performance was not significantly different among treatments (p=0.071, R^2^=0.17, F=2.52, Z=1.43, SS type 1, **Fig 1a, c**), but in *post-hoc* tests, mature workers performed brood care with significantly higher scores than 2-day isolates (x□ = 4.15 and 2.06, respectively; p=0.0093, BH correction). Mature workers scored higher than 40-day isolates (x□ = 1.91; p=0.0093, BH correction). Callows also had a significantly higher brood-care score (x□ score = 3.38) than both 2-day isolates and 40-day isolates (p=0.036 for each pairwise comparison, BH correction). Socially typical callow and mature worker controls were not significantly different, nor were 2-day isolates and 40-day isolates (p = 0.36 and 0.98, respectively).

### Effect of Isolation on Locomotion

Mature controls had significantly higher movement (x□ = 77.84 times a worker moved among quadrants) than workers than 2-day isolates, 40-day isolates, and callows (x□ = 26.38, 38.36, and 18.38 respectively; p=0.004, R^2^=0.35, F=5.97, Z= 2.74, SS type 1, [**Fig 1d, Supp Table Set 1**]); 40-day isolates had the next highest average score (x□ = 38.36). In pairwise comparisons, mature controls were significantly higher than from all other experimental groups (callow, 2, 40: all p<0.0057, BH correction). No other pairwise comparisons were statistically significant (p>0.19).

### Brain Compartment Allometries

Workers experiencing longer periods of isolation had significantly smaller AL volumes compared to those of mature control workers (p=0.007, R^2^=0.39, F=6.19, Z=2.56; **Fig 2a**). Mature workers thus had significantly larger relative AL volumes (on average, 27% relative volume) than 40-day isolates (17% relative volume; p=0.017, BH correction). Forty-day isolates also had significantly lower AL volume than 2-day isolates (23% relative volume, p=0.048, BH correction), suggesting that isolation reduced post-eclosion AL volume. Forty-day isolates had significantly higher relative MB-C volumes (61% relative volume) than control callows (48%) and 2-day isolates (49%; p<0.002, 0.005, respectively; BH correction); MB-C volumes were not significantly different between callows and 2-day isolates (p=0.23; BH correction) or between mature workers (76%) and 40-day isolates (p=0.13, BH correction). The average relative volume of the MB-P showed significant increase between early isolates to older isolates (p=0.029, R^2^=0.21, F=3.74, Z=1.9; **Fig 2b**); however, there was no significant difference in pairwise interactions, indicating that impact on the MB was limited to the calyces, which showed modest growth during isolation (p=0.002, R^2^=0.47, F=9.40, Z=3.52; **Fig 2c**). Other brain compartments (OL, SEZ, CX) did not significantly change with isolation (p=0.25-0.93; **Fig 2d-f**). All additional pairwise analyses of compartmental volumes can be found in **Supp Tables Set 1.**

**Figure 2.**
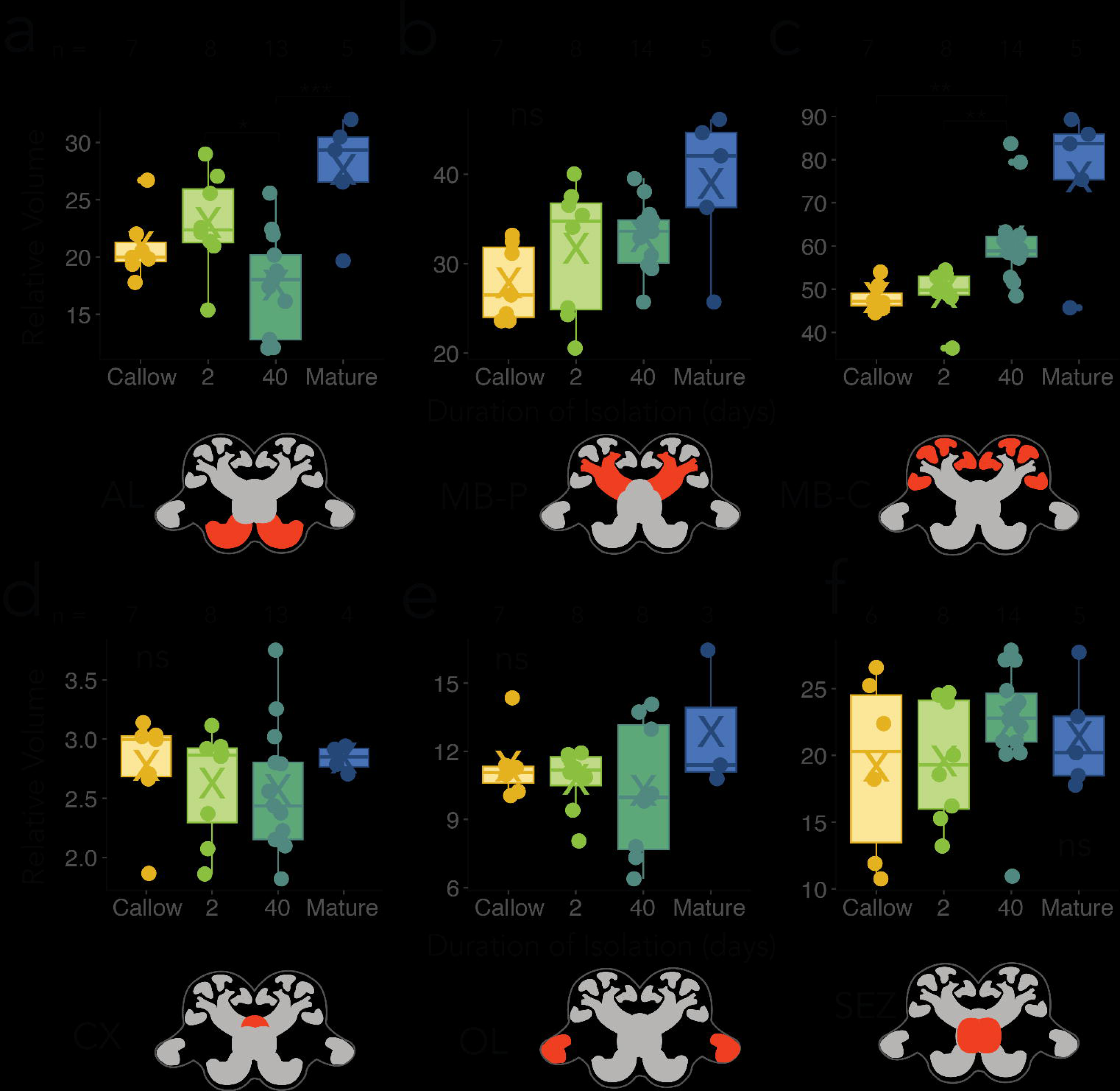
Isolation effects on size of brain compartments. **(a-f)** brain compartment volumes (relative to ROCB) for isolated and control workers. Averages, medians, and samples sizes as in Figure 1. **(a)** AL; **(b)** MB-P; **(c)** MB-C; **(d)** CX; **(e)** OL; **(f)** SEZ. p <0.001 = ***; p<0.001-0.01 = **; p<0.01-0.05 = *; p>0.05 = ns.

### Biogenic amine levels

Brain titers of OA, DA, and 5HT significantly increased with longer periods of isolation (**Fig 3a-c**); levels of 40-day isolates were not significantly different from those of mature workers. DA titers significantly increased with prolonged periods of isolation (p=0.001, R^2^=0.47, F=11.86, Z=3.38, SS type 3, **Fig 3a**) and significantly differed between mature control workers and 2-day isolates, but not between mature controls and 40-day isolates (p < 0.0001, p > 0.05, respectively; BH correction). OA exhibited a similar positive pattern with prolonged periods of isolation (p=0.001, R^2^=0.59, F=32.97, Z=4.96, SS type 3, **Fig 3b**); titers significantly increased between isolated worker groups and with mature workers (p<0.006 for all groups, BH correction). 5HT titers also significantly increased with longer periods of isolation (p=0.001, R^2^=0.61, F=91.77, Z=7.04, SS type 3, **Fig 3c**), with all pairwise comparisons being significantly different (p<0.006) for all groups, BH correction).

**Figure 3.**
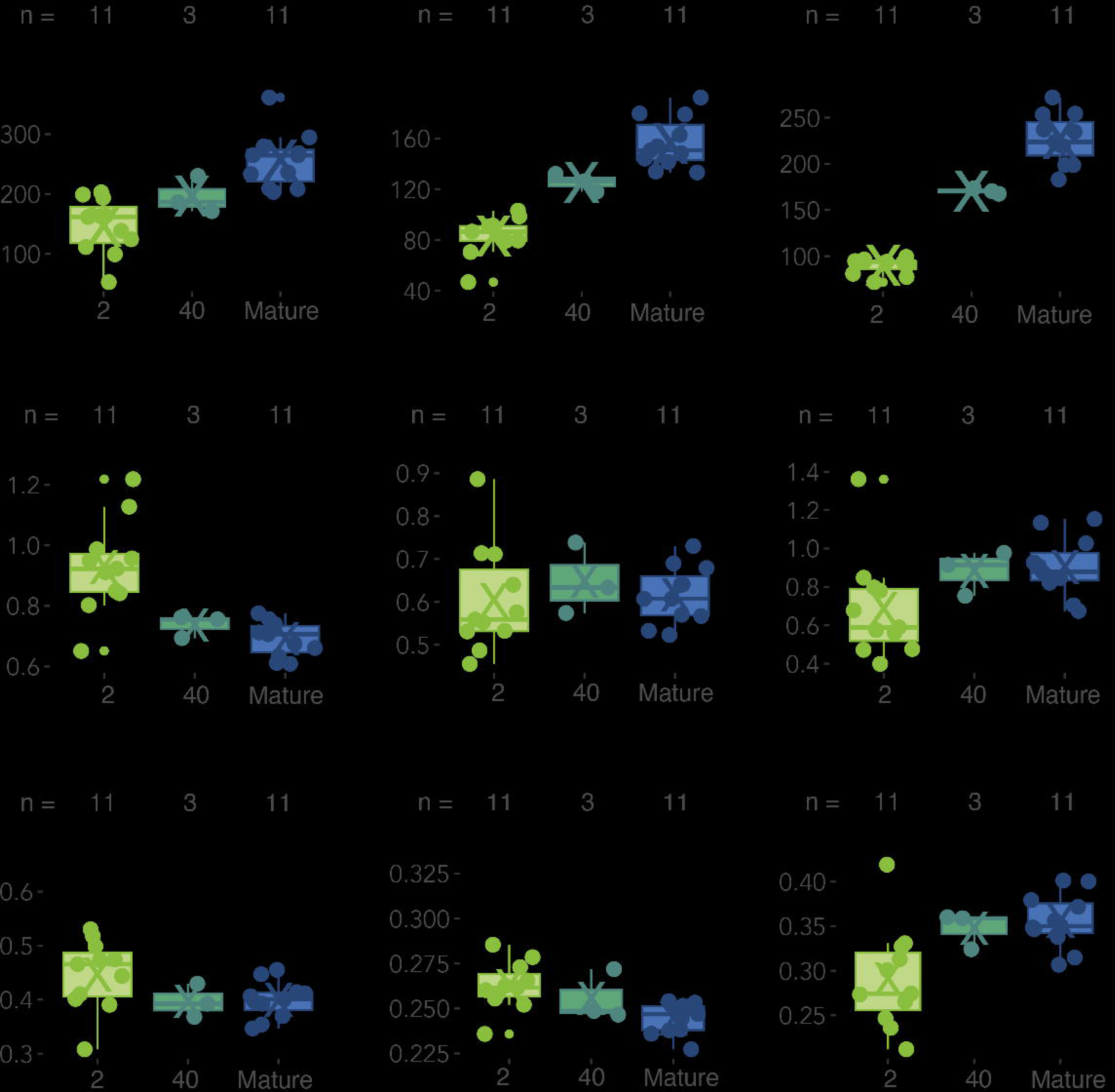
Isolation effects on monoamine levels (a-c), ratios of monoamines (d-e), and their quantities relative to total biogenic amine levels (g-i). Averages, medians, and samples sizes as in Figure 1. p < 0.001 = ***; p<0.001-0.01 = **; p< 0.01-0.05 = *; p>0.05 = ns.

While absolute levels of DA, OA, and 5HT increased from callows to 40-day isolates and matures, their ratios in isolates increased or decreased compared to those recorded in mature controls (**Fig 3d-f**). OA:5HT significantly decreased with time and extended periods of isolation (p=0.004, R^2^=0.21, F=6.37, Z = 3) with significant difference between 2-day isolates and mature controls (p < 0.001, BH correction; **Fig 3d**). Forty-day isolates were not significantly different from any treatment category. Neither OA:DA (p=0.96, R^2^=0.0041, F=0.048, Z= −1.74) nor 5HT:DA (p=0.40, R^2^=0.061, F=1.00 Z=0.30) significantly changed with isolation. However, 2-day isolates had significantly lower 5HT:DA ratios than mature workers (p=0.02, respectively; BH correction).

DA titers relative to total monoamine levels did not significantly differ across isolation treatment groups and mature workers (p=0.47, R^2^=0.050, F=0.77, Z=0.057). Relative OA levels, in contrast, were significantly higher in younger isolates compared to mature workers (p=0.034, R^2^=0.26, F=4.14, Z=1.79); 2-day and mature controls were significantly different (p=0.002, respectively, BH correction). Relative 5HT quantities did not change in isolation treatments and in comparison, to mature controls (p=0.40, R^2^=0.061, F=1.00 Z=0.30). However, 2-day isolates had significantly lower 5HT:DA ratios compared to 20-day isolates and mature controls (p=0.02, respectively; BH correction). Relative 5HT quantities were significantly different (p=0.094, R^2^=0.13, F=3.65, Z=1.32) between 2-day vs. mature controls (p<0.008 respectively, BH correction). All additional pairwise analyses for biogenic amine levels can be found in **Supp Tables, Set 1.**

## Discussion

Brood-care and predatory behavior scores of *C. floridanus* workers isolated for varying durations after eclosion were lower than those of control workers that developed in a natural social environment. Two-day isolates showed a similar low level of predatory behavior as that of callow controls, but the predation scores of 40-day isolates were significantly lower than those of mature control workers, suggesting impairment due to isolation. Social isolation also resulted in a significant difference in pupal care – as measured in our assay – between 2-day isolates and callow control workers, and 40-day isolates and mature control workers. This result may reflect the development of behavioral competencies associated with myological maturation [54] that may be compromised by a lack of sensory inputs. Workers in all isolation treatment groups showed significantly decreased locomotion relative to mature workers **(Fig 1d, Supp Fig 1c**), consistent with Wang et al. [23] for isolated bumblebees but different from *Cataglyphis niger* and *Camponotus fellah* ant workers, which increased movement following isolation and then decreased movement [22, 55]. Older isolates (40-day isolates) generally scored higher than workers isolated for a shorter period (2-day isolates), but were still significantly lower in their activity, predatory, and brood-care scores than mature controls. This suggests that some behaviors develop in an experience-expectant manner and require social contact for complete development. *C. floridanus* may thus exhibit a combination of experience-expectant and experience-dependent components. The latter has been found in honeybees, which show an increase in mushroom body size with foraging experience [29].

*C. floridanus* worker brains normally increase in size during a 10-month period after adult eclosion, the greatest growth rate (nearly 100%) occurring in the MBs and ALs [28]. Our maximum early life isolation treatments were approximately 12-17% of this period of maturation. Workers isolated for longer durations and socially typical mature workers generally had larger relative MB-C volumes compared to workers isolated for shorter periods and socially typical callows. MB-C size in 40-day isolates was not significantly different from that of mature controls, suggesting gross neuroanatomical development remained stable under conditions of isolation. These findings differ from the results of Seid and Junge [17], which showed that MBs of isolated *C. floridanus* workers do not change in volume relative to total brain size. MB-C resilience through 71 days of isolation has been found in paper wasps [56]. Using bumblebees as a model, Wang et al. [23] associated expression differences in juvenile hormone-related genes and transcriptomic network dysregulation with isolation-induced variation in brain volume.

Bumblebees isolated after eclosion for 9 days showed significant differences in the variance of MB volume rather than average volume compared to socially typical bees, suggesting growth destabilization in some, but not all, workers. On the other hand, our results agree with research showing that the mushroom bodies of isolated honeybees continue to grow [57]. Our study identified differential sensitivity and resilience in brain compartment growth. Although we did not assess temporal patterns to determine if the MBs or other compartments accelerated or decreased differentially in size over time in isolates and/or socially typical workers, results nevertheless suggest that MB development is resilient to social isolation.

In contrast, AL volume decreased with periods of isolation approximately 5 days or longer; 40-day isolates had, on average, smaller ALs than control callows and mature workers. This decline in AL size is similar to the findings of Wang et al. [23] on bumblebees. Because AL size decreased with isolation duration in *C. floridanus,* loss of olfactory ability could decrease brood-care and predatory behavior by 40-day isolates compared to mature controls. Social contact early in life thus appears to provide sensory inputs necessary to stimulate a level of AL development required to perform the tasks we assayed. Isolation did not impact OL volume in our study of *C*. *floridanus*, although it did reduce OL volume in paper wasps, which integrate visual information used in conspecific facial recognition [56, 58]. The CX, which functions in the control of movement [59] and sensorimotor processing in insects [60, 61], did not change significantly in volume in isolated *C. floridanus* workers, a result inconsistent with Wang et al. [23] for isolated bumblebees. The volume of the SEZ, which regulates mouthparts [62] and is likely involved in the control of brood-care motor functions, did not significantly change with isolation. This supports that the decline in brood-care behavior may have been due to a deficit in olfactory processing rather than mouthpart motor deficits.

Although it may be generally assumed that smaller neuropil size reduces competence, we do not know how whole-brain or compartmental volumes affect behavioral ability. Indeed, the relationship between brain size and behavioral capacity is actively debated [72]. In ants, elaboration of the MB appears to represent increased demands for higher-order processing [26, 73, 74] and larger OLs are correlated with more complex visual ecologies [75,76]. Larger ALs have been associated with age-related olfactory ability [64]. However, Hart et al. [77] showed that ant alarm pheromone response is encoded by less than 10 of a total of approximately 500 AL glomeruli, a result suggesting that the relationship of neuropil size and behavior is unclear. We also do not understand how circuit configuration, activation, synaptic strength, or other neural processes in the ant brain connectome are influenced by worker isolation.

Worker behavioral maturation, including repertoire expansion and a corresponding increase in brain size, correlates with increasing monoamine levels and the development of serotonergic circuitry in some ants [32, 63–65]. Our *C. floridanus* data indicate that levels of OA, DA, and 5HT increase with age in the brains of isolated workers, despite a lack of social contact. Indeed, 40-day isolates had monoamine levels like those of mature workers. Ratios of biogenic amines can significantly change with age in ants and may influence the development of age-related behavior [32] by modifying response thresholds. However, we found no significant differences in neuromodulator ratios between 40-day isolates and mature controls, indicating that aminergic circuitry may be maintained under conditions of social deprivation. Boulay et al. [30] and Wada-Katsumata [47] found that 5HT and DA levels were robust to social isolation and Boulay et al. [30] hypothesized OA may reduce the effects of social deprivation, acting homeostatically to maintain social cohesion. Whether octopaminergic signaling has a stimulatory or inhibitory effect on prosocial behaviors such as trophallaxis, which may be affected by periods of social isolation, requires further study.

Neuromodulators can restore sensorimotor functions following injury [72], alter synaptic functions to significantly change circuitry [73, 74], and increase the robustness of network functions [75] to recover from disruption and/or stress [76, 77]. In the ant *Pheidole dentata*, monoamine levels continue to increase in titer in aging workers, which do not appear to decline in task performance capability [38]. Given that DA, OA, and 5HT can flexibly reshape circuits, neuromodulators may reduce isolation-induced neuroanatomical deficits in ants, stabilizing circuit dynamics or returning them to a stable state if disrupted. Although we did not identify an increase in monoamine levels in isolates, this does not preclude neuromodulatory compensation to maintain circuitry and behavioral functions under conditions of social deprivation.

## Conclusion

The results of prior research on the impacts of senescence on neurochemistry, neuroanatomy, apoptosis, and behavior [38], as well as deprivation of experience [20] and reduced olfactory processing ability [39], suggest that ant brains can have the capacity to buffer insults of aging and physical damage to sensory appendages. The present study suggests brain compartments vary in susceptibility and resilience to social deprivation and that aminergic modulatory systems are robust to isolation. The 36% mortality of isolated *C. floridanus* workers we recorded indicates a significant negative effect that may be due more to social processes affecting nutrition and energy availability [22, 55] than neurobiological deficits resulting from isolation. Workers that survive neurobiological and physiological “bottlenecks” appear able to recover from challenges to some processes of brain development, which in *C. floridanus* are dependent on age as well as experience [28]. Age-related increases in brain size in ants appear to be due to stable programmed axonal and dendritic growth processes. If sensory experience is lacking or temporally atypical, brain development may still proceed in part in the absence of inputs or may be delayed (e.g., [78]). In our study, the effect of social isolation in *C. floridanus* appears to reflect such robust programming, perhaps due in part to stabilizing effects of neuromodulators.

The resilience of worker ants to challenges during development may functionally resemble differences that enable some individuals – perhaps those with adaptive synaptic phenotypes [79] or ion channel properties [80] to withstand neural damage and preserve behavior. We hypothesize that the ability to buffer stressors may involve such variation in circuitry that characterizes clade-wide neuroarchitectures of the miniaturized brains of ants, acknowledging that few species have been studied. Analyses of temporal patterns of brain gene expression in *C. floridanus* isolated workers can identify transcriptomic changes linked to brain development and influenced by social experience [81, 82] and possibly identify genetic mechanisms of resilience. Genomic studies of phylogenetically and sociobiologically diverse ant species can also contribute to our understanding of how workers are susceptible to and/or may recover from social deprivation or are resilient to isolation.

## Supporting information

Isolation Supplement

Supplementary Tables, Set 1

Supplementary Tables, Set 2

## Acknowledgements

We thank Lloyd Davis for colony collection. BCG thanks Dr. Katherine Fiocca for manuscript feedback and Dr. Bryan Juarez for coding advice. We thank two anonymous reviewers for their thoughtful comments.

## Statement of Ethics

Ant colonies were cultured in an environment for optimal growth. Workers were cold anaesthetized before dissection. Other issues of compliance with animal care regulations do not apply. An ethics statement was not required for this study, as no humans nor vertebrates were used in accordance with Boston University Institutional Animal Care and Use Committee.

## Author Contributions Statement

BCG, IBM, and JFAT designed the study. BCG recorded behavior. BCG and IBM immunohistologically prepared and imaged brains. EJS quantified brain volume. BCG, ZC, and MM measured monoamine titers. BCG statistically analyzed volumetric data. BCG drafted the manuscript. BCG, EJS, IBM, ZC, MM, and JFAT developed and edited the manuscript.

---

**Supplementary Figure 1.**
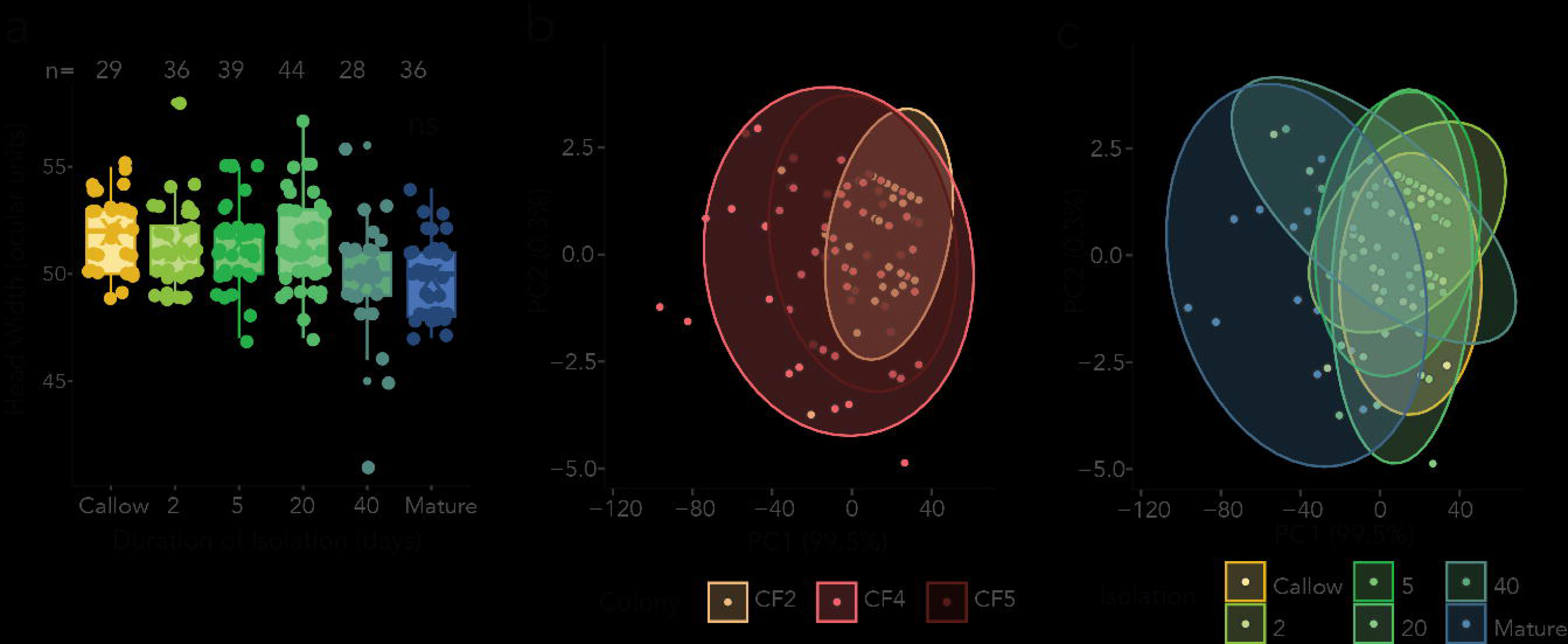
**(a)** Head width variation of sampled workers **(B-C)** Principal component analyses of behavior by **(b)** colony identity and **(c)** isolation duration. Colony identity and isolation duration do not have significant effects. Significance level: <0.001 = ***; 0.001-0.01 = **; 0.01-0.05 = *; >0.05 = ns. X = average; horizontal line = median. ns = not significant.

**Supplementary Figure 2.**
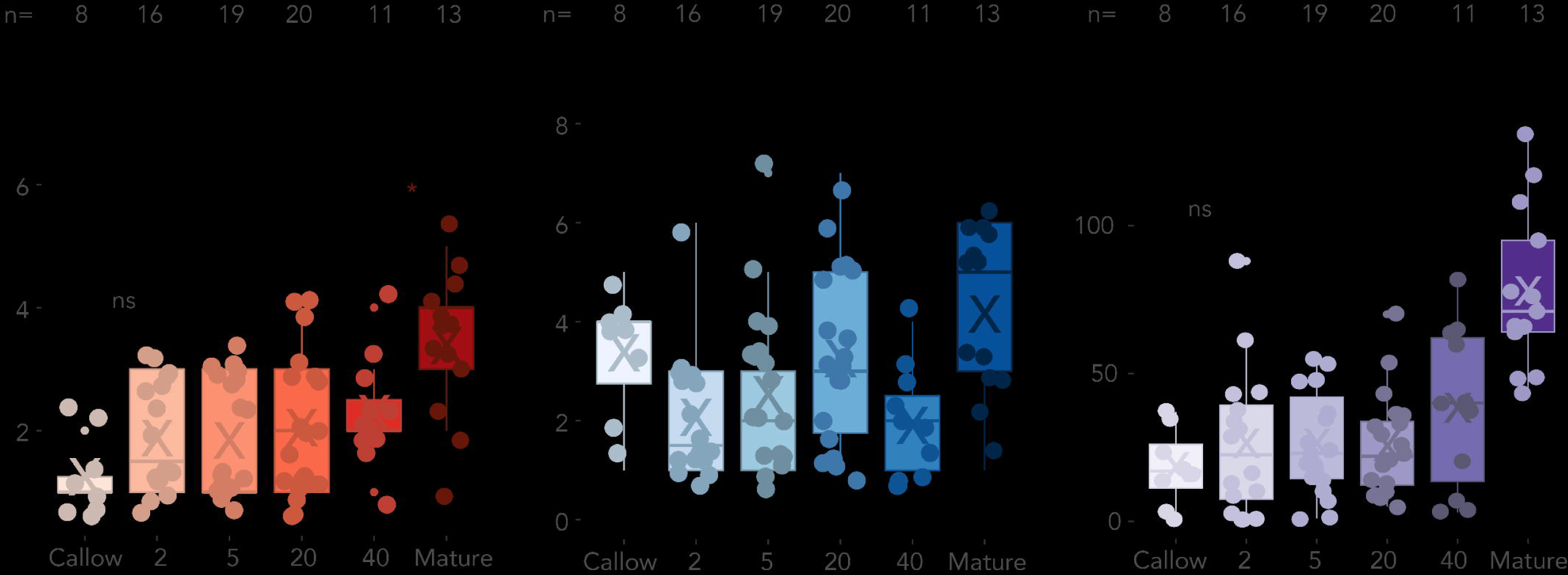
Task performance trajectories of isolated and control workers. **(a)** Illustration of assays to assess brood-care and predatory behaviors. **(b)** Predation scores of isolated and control workers. **(c)** Brood-care scores of isolates, callows, and mature worker controls. **(d)** Locomotion scores of isolates, callows and mature controls. **(b)**-**(d)**, X = average; horizontal line = median. Sample sizes are above box plots. <0.001 = ***; 0.001-0.01 = **; 0.01-0.05 = *; >0.05 = ns.

**Supplementary Figure 3.**
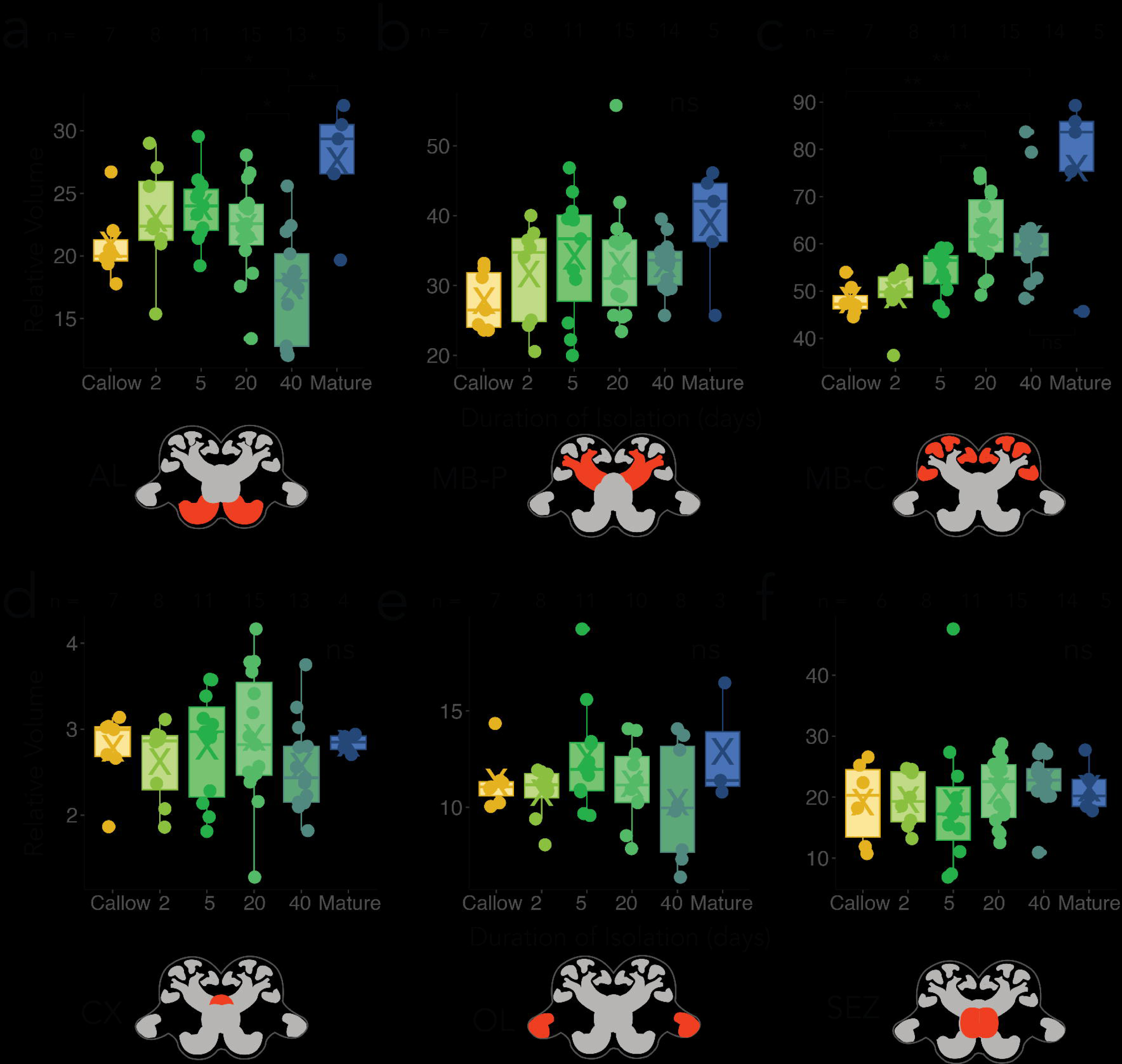
Brain compartment growth trajectories for isolated workers. **a-e**: brain compartment volumes (relative to ROCB) for isolated and control workers. Sample sizes of each group are above box plots. X in each box plot represents the average and horizontal line the median of each group **(a)** AL; **(b)** MB-P; **(c)** MB-C; **(d)** CX; **(e)** OL; **(f)** SEZ. p <0.001 = ***; p<0.001-0.01 = **; p<0.01-0.05 = *; p>0.05 = ns.

**Supplementary Figure 4.**
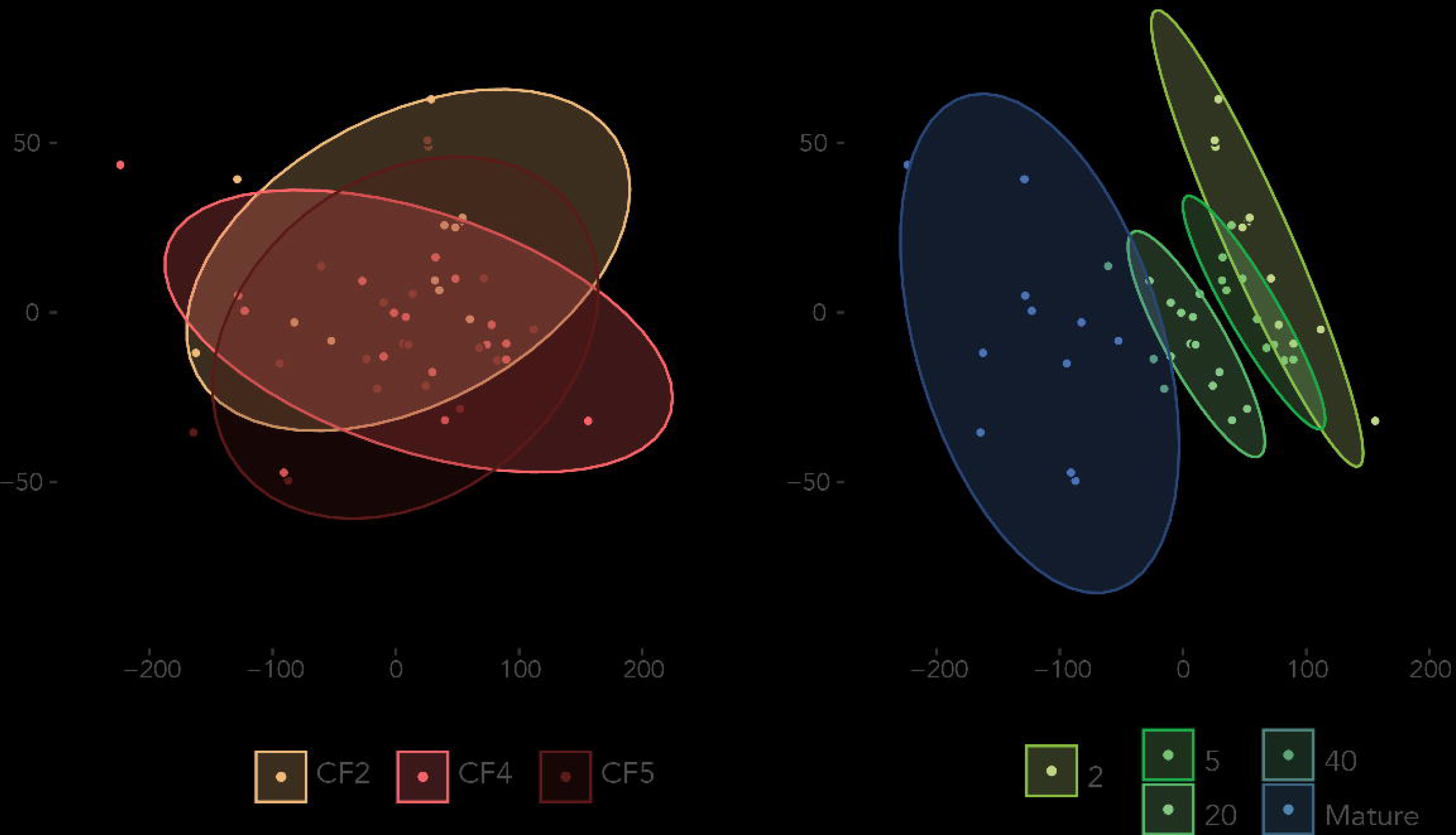
Trajectories and colony effects on biogenic monoamine levels. **(a-b)** Principal component analyses of biogenic amines clustered by (**a**) colony identity and (**b**) isolation duration.

**Supplementary Figure 5.**
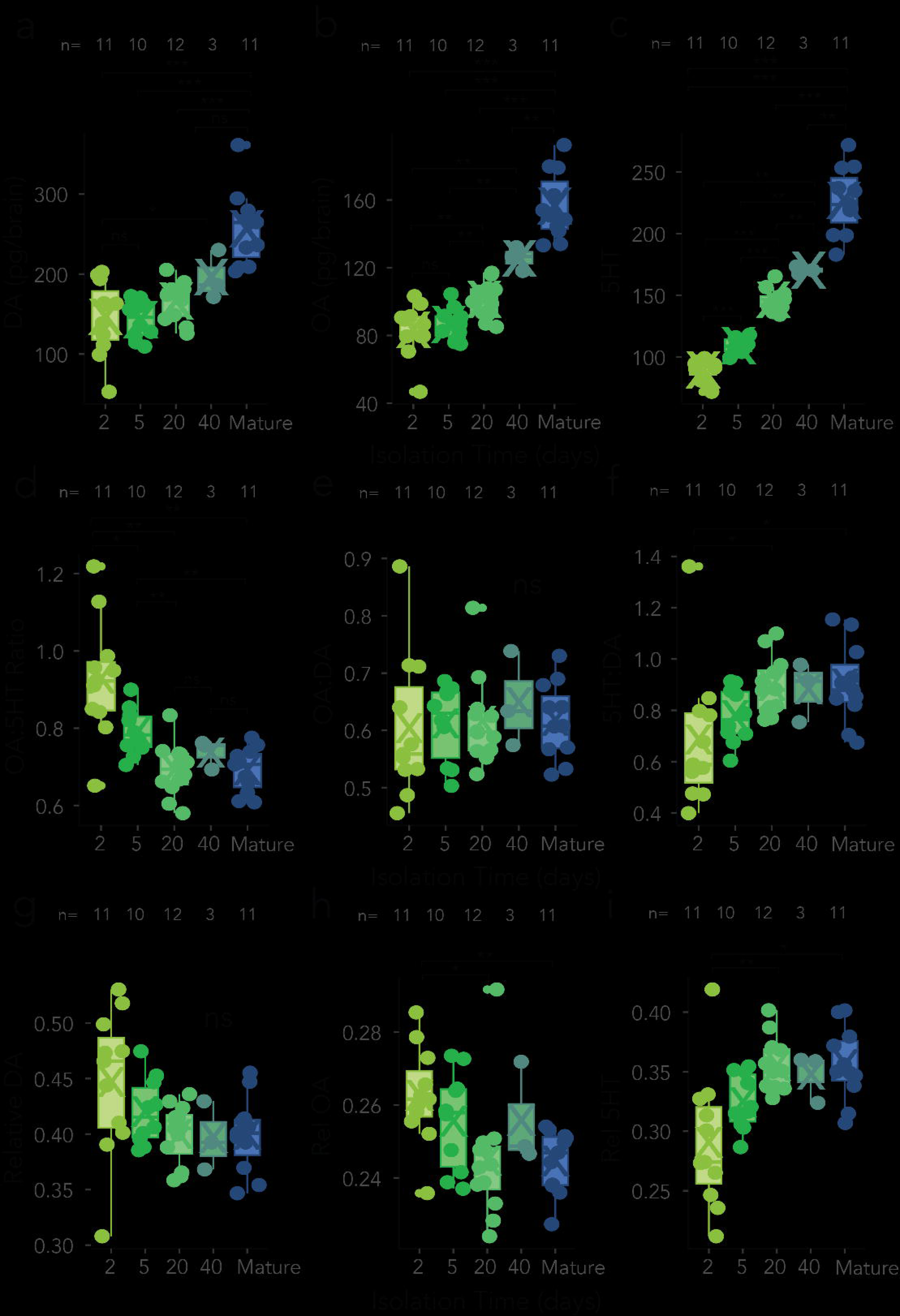
Biogenic amine levels across isolation trajectories (a-c), ratios of DA, OA, and 5HT (d-f), and levels of DA, OA, and 5HT to total biogenic amines (g-i). **(c-e)** Trajectories of biogenic amine titers in response to isolation period. (d-f) Trajectories of biogenic amine titer ratios in response to isolation period. (h-i) Relative biogenic amine to total amine titer ratios. Averages, medians, and samples sizes as in Figure 3. p <0.001 = ***; p<0.001-0.01 = **; p< 0.01-0.05 = *; p>0.05 = ns.

