## Supplementary material for "Differential Neuroanatomical, Neurochemical, and Behavioral Impacts of Early-Age Isolation in a Eusocial Insect": Isolation Supplement

***Supplementary Information***

***Supplementary Data***

Supplementary raw data are available for download in Supplementary files brainvolumetricdata.xlsx, behaviorandheadwidth.xls, and biogenicamines.xlsx

Supplementary tables summarizing statistical details **excluding** 5- and 20-day isolates are available for download in Supplementary file SupplementaryTables_set1.xlsx. For statistical analyses **including** 5- and 20- day isolates, please see Supplementary file SupplementaryTables_set2.xlsx.
